# Limited effects of m^6^A modification on mRNA partitioning into stress granules

**DOI:** 10.1101/2021.03.19.436090

**Authors:** Anthony Khong, Tyler Matheny, Thao Ngoc Huynh, Vincent Babl, Roy Parker

## Abstract

Recent studies have argued that the m^6^A modification of mRNAs promotes mRNA recruitment to stress granules through the interaction with YTHDF proteins (Anders et al., 2018; Ries et al., 2019). However, mRNAs that contain multiple m^6^A modified sites partition similarly into stress granules in both wild-type and m^6^A-deficient cells by single-molecule FISH suggesting m^6^A modifications play a minor role in mRNA partitioning into stress granules. Moreover, multiple linear regression analysis suggests m^6^A modification plays a minimal role in stress granule recruitment. Finally, the artificial tethering of 25 YTHDF proteins on reporter mRNAs leads to only a modest increase in mRNA partitioning to stress granules. These results indicate m^6^A modification makes a small, but measurable, contribution to recruiting specific mRNAs to stress granules.

## INTRODUCTION

Stress granules are cytoplasmic molecular condensates composed of non-translating messenger ribonucleoproteins (mRNPs). Stress granules form when there is an increase in the pool of non-translating mRNPs, which often occurs when cells undergo a variety of stress conditions including oxidative stress, hypoxia, and heat shock that downregulate bulk translation (Ivanov et al., 2019; Protter & Parker, 2016). Stress granules are of interest since they are thought to play roles in a variety of diseases such as viral infection, cancer, and neurodegenerative disorders. Moreover, their investigation may provide insights into other RNA and proteins condensates such as the nucleolus, P-bodies, and germ granules (Lloyd, 2013; Protter & Parker, 2016; Song & Grabocka, 2020; Wolozin & Ivanov, 2019).

Recent studies have elucidated the composition of stress granules in a variety of stress conditions (Jain et al., 2016; Khong et al., 2017; Markmiller et al., 2018; Namkoong et al., 2018; Padrón et al., 2019; Youn et al., 2018). One of the major conclusions from these studies is the mRNAs that are preferentially enriched in stress granules are biased towards poorer translation efficiency and longer length (Khong et al., 2017; Namkoong et al., 2018). Despite this length difference, mRNAs of the same length can show different recruitment into stress granules arguing that there can also be sequence-specific information that affects mRNPs partitioning into stress granules. Indeed, recent studies have concluded that the m^6^A modification of mRNAs can increase their partitioning into stress granules by the binding of YTHDF proteins (Anders et al., 2018; Ries et al., 2019). However, no study directly tested this conclusion by inhibiting m^6^A modification and examining the effect on mRNP partitioning. Thus, we compared the localization of poly-m^6^A mRNAs in wildtype and ΔMETTL3 mES cells in stress granules and discovered the poly-m^6^A mRNAs are enriched similarly in stress granules in both cell types. These results suggest m^6^A plays a small role in recruiting RNA to stress granules. Moreover, we observed tethering up to 25 YTHDF proteins to a reporter mRNA has only a modest increase in the reporter mRNA partitioning into stress granules. These results suggest individual m^6^A-YTHDF interactions play a small role in recruiting RNAs to stress granules.

## RESULTS AND DISCUSSION

In previous work, three main observations have been presented to argue that m^6^A modification targets mRNAs to stress granules by providing binding sites for YTHDF proteins, whose intrinsically disordered regions (IDRs) then promote mRNPs entering stress granules (Reis et al., 2019). First, they show YTHDF proteins can undergo self-assembly *in vitro* through liquid-liquid phase separation (LLPS), which is dependent on their IDRs, and increased by m^6^A modified RNAs. However, whether this *in vitro* LLPS is relevant to the cell was not tested. This is an issue since many proteins can undergo LLPS *in vitro*, particularly at high concentrations and in the absence of competing proteins (Protter et al., 2018).

A second observation used to argue m^6^A modification promotes mRNAs entering RNP granules is the demonstration that the YTHDF2 protein localizes to P-bodies in the absence of stress and to stress granules during stress. However, the partitioning of YTHDF2 is dependent on RNA binding since YTHDF2 fail to associate with stress granules or P-bodies when RNA binding is blocked by deletion of the m^6^A methylase (ΔMETTL14), or by expressing an RNA-binding mutant YTHDF2 protein in cells. This argues that YTHDF2 partitioning into RNP granules is due to binding to RNA and indicates that the YTHDF2 IDR is not sufficient for stress granule partitioning. The simplest interpretation of this observation is that when YTHDF proteins are bound to mRNAs, they can be carried into mRNP granules such as P-bodies and stress granules.

The third argument presented is that m^6^A strongly promotes the recruitment of poly-m^6^A mRNA to stress granules relies on analyzing individual mRNA localization to stress granules by single-molecule FISH. Specifically, it is reported that two poly-m^6^A mRNAs, *Fem1b and Fignl1*, are recruited to stress granules more effectively (~60%) than two non-methylated m^6^A mRNAs, *Grk6* and *Polr2a*, to stress granules (~30%). Since these mRNAs are of similar size, which is relevant since length can affect mRNAs partitioning into stress granules, it was inferred that their difference in stress granule recruitment was due to differential m^6^A methylation.

In principle, the differential localization of the *Fem1b, Fignl1, Grk6*, and *Polr2a* mRNAs could be due to m^6^A differences or to other features of each mRNA. To directly test if m^6^A is affecting the localization of mRNAs into stress granules, we analyzed the partitioning of *Fem1b* and *Fignl1* mRNAs into stress granules in wildtype and ΔMETTL3 mES cells (Batista et al., 2014), which removes the m^6^A mRNA writer (Liu et al., 2014) and its knockout reduces m^6^A levels on mRNAs in mES cells (Figure 1A).

**Figure 1.**
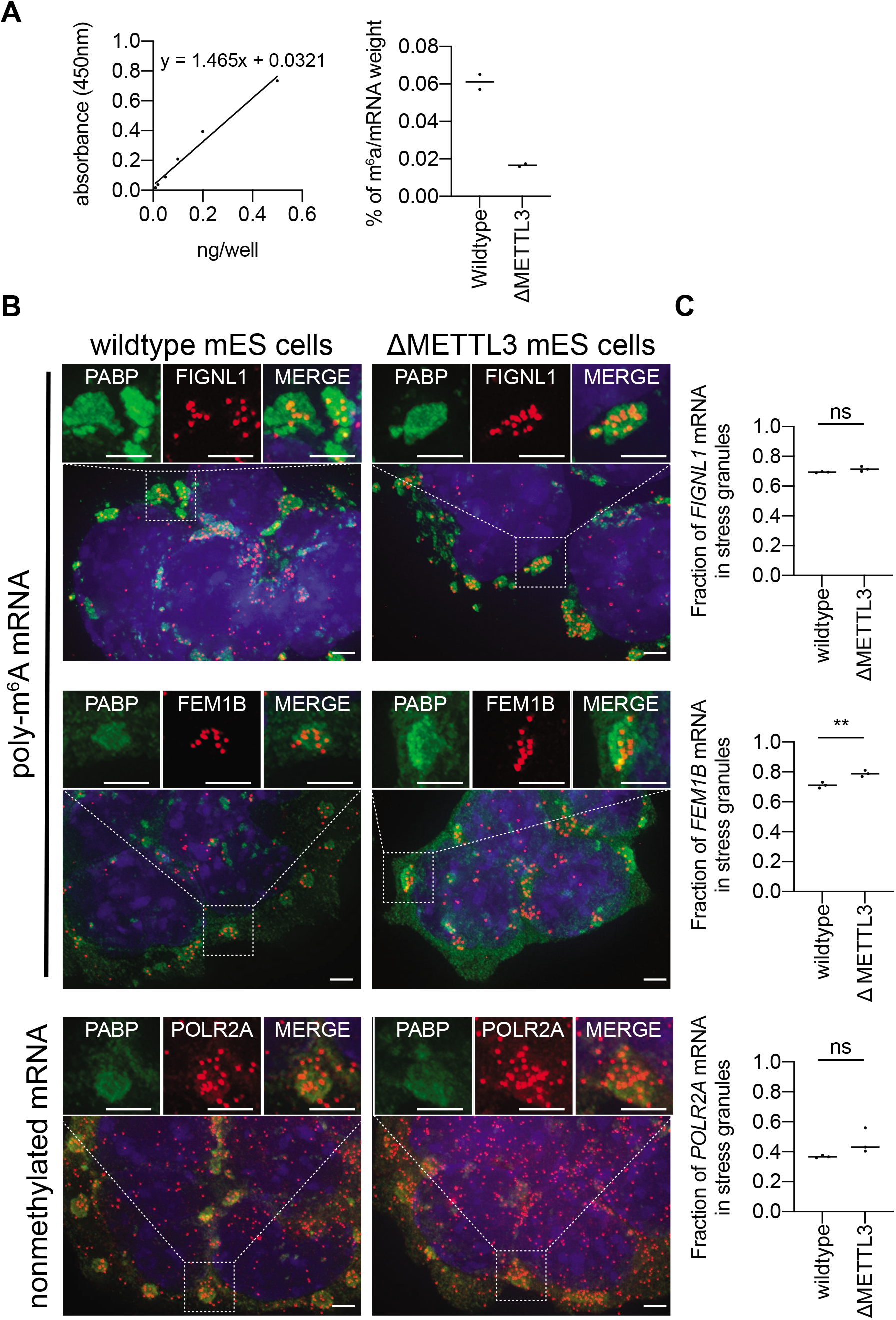
There is no difference in the fraction of polymethylated m^6^A mRNAs in arsenite-induced stress granules between wildtype and ΔMETTL3 mES cells.

(**A**) Left graph: OD450 absorbance values from m^6^A standard controls provided by EpiQuick m^6^A RNA Methylation Quantification Kit was measured and a linear regression was plotted in order to quantify the absolute amount of m^6^A in mRNA isolated from wildtype and ΔMETTL3 mES cells shown in the right graph. Right graph: Fraction of m^6^A weight relative to all nucleic acid weight in mRNAs between wildtype and ΔMETTL3 mES cells. (**B**) Representative images of wildtype and ΔMETTL3 mES cells stressed for one hour with arsenite and costained with single-molecule FISH probes against Fignl1, Fem1b, and Polr2A (red) and antibody against G3BP1 protein (green). The nuclei are stained with DAPI (blue). The scale bar is 3 μm. (**C**) Scatter plot of the fraction of RNA molecules in stress granules in U-2 OS and ΔMETTL3 U-2 OS cells. Three biological replicates were performed. ** and ns denote P < 0.01 and not significant respectively.

Strikingly, we observed no significant differences between WT and ΔMETTL3 mES cells in the stress granule accumulation of the *Fem1b* and *Fignl1* mRNAs (Figure 1B, C), each of which contain 4 mapped m^6^A sites (Xiang et al., 2017). These results argue m^6^A modification does not play a major role in the targeting of these mRNAs to stress granules.

These results indicate individual m^6^A interaction with YTHDF protein is not a strong contributor to mRNA partitioning in stress granules or does not play a role at all. However, prior work has described a correlation between the number of m^6^A sites on mRNAs (Ries et al., 2019; Xiang et al., 2017) and enrichment in stress granules or mRNP granules in Khong et al., (2017) and Namkoong et al., (2018), respectively. However, this correlation may be fortuitous since both the number of m^6^A sites and stress granule enrichment are correlated with mRNA length (Figure 2A) (Khong et al., 2017).

**Figure 2:**
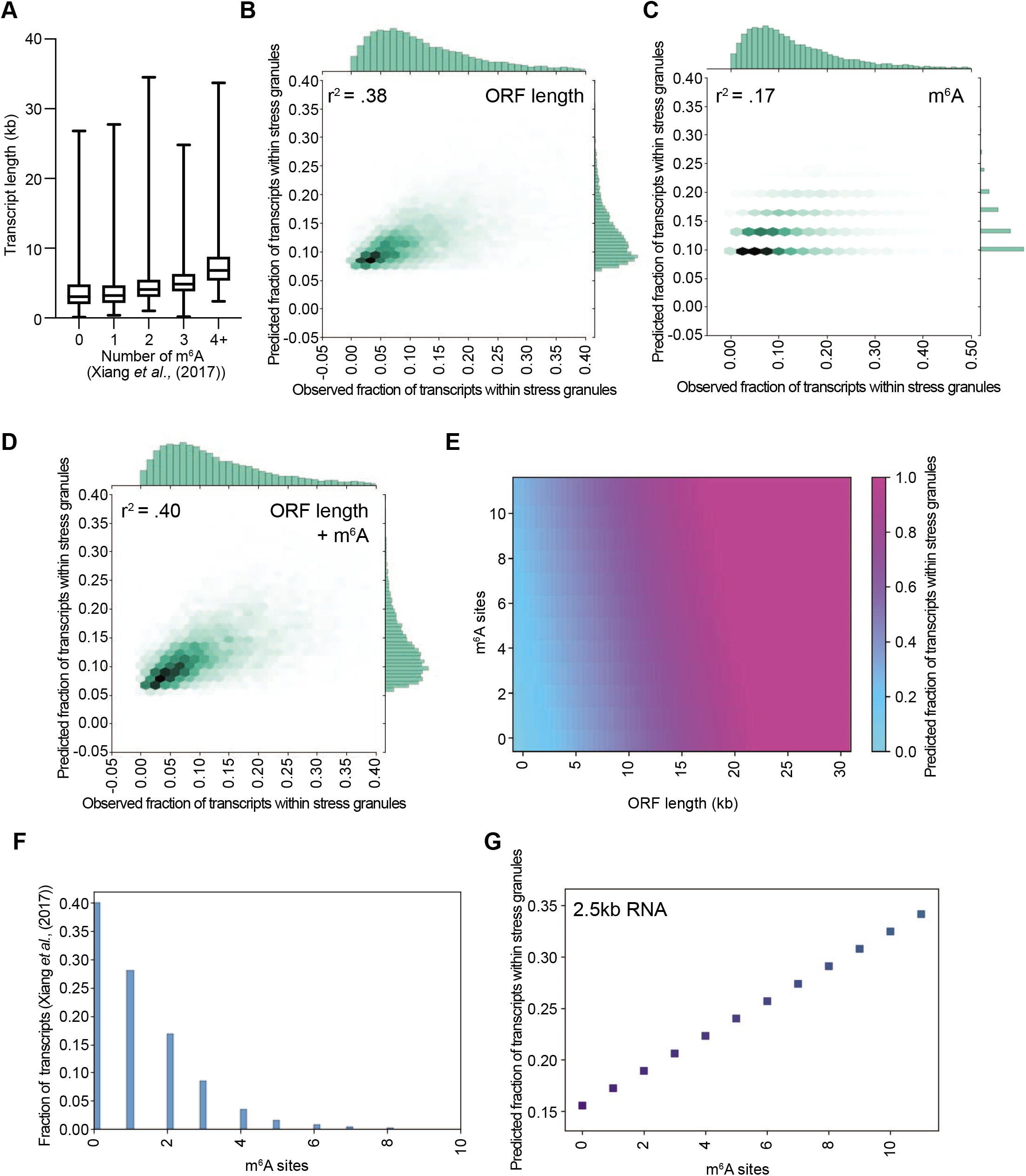
Bioinformatic prediction of m^6^A contribution to stress granule enrichment.

(**A**) Box plot of transcript length versus the number of mapped m^6^A sites in U-2 OS cells (Xiang *et al.*, 2017) (B) Scatterplot depicting predicted vs observed fraction of transcripts within stress granules. Predicted values were obtained from the linear regression model was based on ORF length alone. (C) Same as A, but using a linear model based on the number of m^6^A sites per transcript. (D) Same as A, but using a linear model based on both ORF length and number of m^6^A sites per transcript. (E) Heatmap showing predicted fraction of transcripts within stress granules for simulated transcripts of various lengths and m^6^A levels. (F) Histogram depicting the fraction of transcripts containing various m^6^A sites. (G) Scatterplot depicting predicted fraction of transcripts within stress granules vs the number of m^6^A sites.

To computationally examine if m^6^A modification contributes to mRNA partitioning into stress granules independent of length, we performed multiple regression analyses where we compared the effect of mRNA ORF length on stress granule partitioning with or without an additional contribution of m^6^A modification. We found that a linear regression model based on ORF length alone showed an R^2^ score of .38 for predicted stress granule enrichment vs observed stress granule enrichment (Figure 2B). A linear regression model based on the total number of m^6^a sites per transcript showed an R^2^ score of .17 for predicted stress granule enrichment vs observed stress granule enrichment (Figure 2C).

While ORF length was a better predictor of stress granule enrichment than m^6^A modification, this does not rule out the possibility that m^6^A plays an additional role in stress granule enrichment in concert with length. To parse the relative contributions of length and m^6^A to stress granule enrichment, we built a multiple linear regression model using length and m^6^A as predictors of stress granule enrichment (Figure 2 and Supplemental Figure 1). When we considered the combination of m^6^A and ORF length, our model improved from an R^2^ value of .38 to .40, suggesting that m^6^A explain an additional 2% of the variance in stress granule enrichment (Figure 2D). We see similar results when we considered the combination of m^6^A and overall length (Supplemental Figure 1). We also see nearly identical results where m^6^A provides a small increase in explaining the variance in stress granule enrichment using another m^6^A metric, m^6^A ratio (Molinie et al., 2016) (Supplemental Figure 1). These results suggest that m^6^A modification provides a minimal impact on mRNA partitioning into stress granules.

While the addition of m^6^A modifications improved our R^2^ metric from .38 to .40, the R^2^ metric does not give any insight into to what degree m^6^A might enhance stress granule enrichment. To examine the degree to which ORF length and number of m^6^a sites contribute to stress granule enrichment, we simulated data representing transcripts of lengths up to 30 kb and ranging from 0 to 11 m^6^A sites and analyzed the predicted stress granule enrichment values of the multiple linear regression model (Figure 2E). We observed that transcript length showed a stronger influence on stress granule enrichment than m^6^A modifications (Figure 2E), which can be visualized by the striking color change across the ORF length axis (x-axis). The number of m^6^A modifications did show some effect on stress granule enrichment (y-axis), and we note that as the number of m^6^A sites per transcript increase (y-axis), stress granule enrichment also is predicted to slightly increase.

It should be noted that this simulation is likely overestimating the contributions of m6A modifications for two reasons. First, we simulated transcripts containing up to 11 m^6^A sites, but >95% of transcripts are thought to have 4 or fewer m^6^A sites (Figure 2F) (Xiang et al., 2017). Second, in this analysis we assume that each m^6^A site is 100% modified, which is unlikely to be true since often specific m^6^A modification sites are modified at lower rates (Liu et al., 2013).

In order to examine the degree to which we would expect m^6^A modifications to influence stress granule partitioning, we plotted predicted stress granule enrichment as a function of the number of m^6^A sites for an RNA of a fixed 2.5 kb ORF length. Even assuming 100% modification of each site, we found that each additional m^6^A modification on an RNA of a fixed 2.5 kb length would lead to a ~1.7% increase in stress granule enrichment (Figure 2G). Thus, for the majority (>95%) of transcripts, which contain 4 or fewer m^6^A sites, we would predict m^6^A to maximally account for a 6.8% increase in enrichment.

Our multiple linear regression model is limited by the fact that we compare m^6^A and stress granule enrichment from two different studies (Khong et al., 2017; Xiang et al., 2017). Thus, we sought to directly test our model by other means. To test the merit of our linear regression model and experimentally quantify how much YTHDF-m^6^A interaction plays a role in recruiting RNA stress granules, we tethered 25 YTHDF proteins on luciferase reporter RNAs and examine its localization to stress granules using the λN-BoxB system (Matheny et al., 2021). YTHDF1 and 2 fused to GFP-λN transgene are transfected into U-2 OS cells stably expressing a luciferase reporter containing 25-BoxB stem-loops (Figure 3A). We observed the tethering of YTHDF1-GFP-λN and YTHDF2-GFP-λN are functional because their expression reduced 25-BoxB-luciferase reporter levels, but not 0-BoxB-luciferase reporter levels (Figure 3B), which is consistent with these YTHDF proteins enhancing mRNA degradation when associated with mRNAs (Zaccara & Jaffrey, 2020). We then stressed the cells with arsenite for 60 minutes and quantified the localization of the reporter mRNA to stress granules by single-molecule FISH.

**Figure 3.**
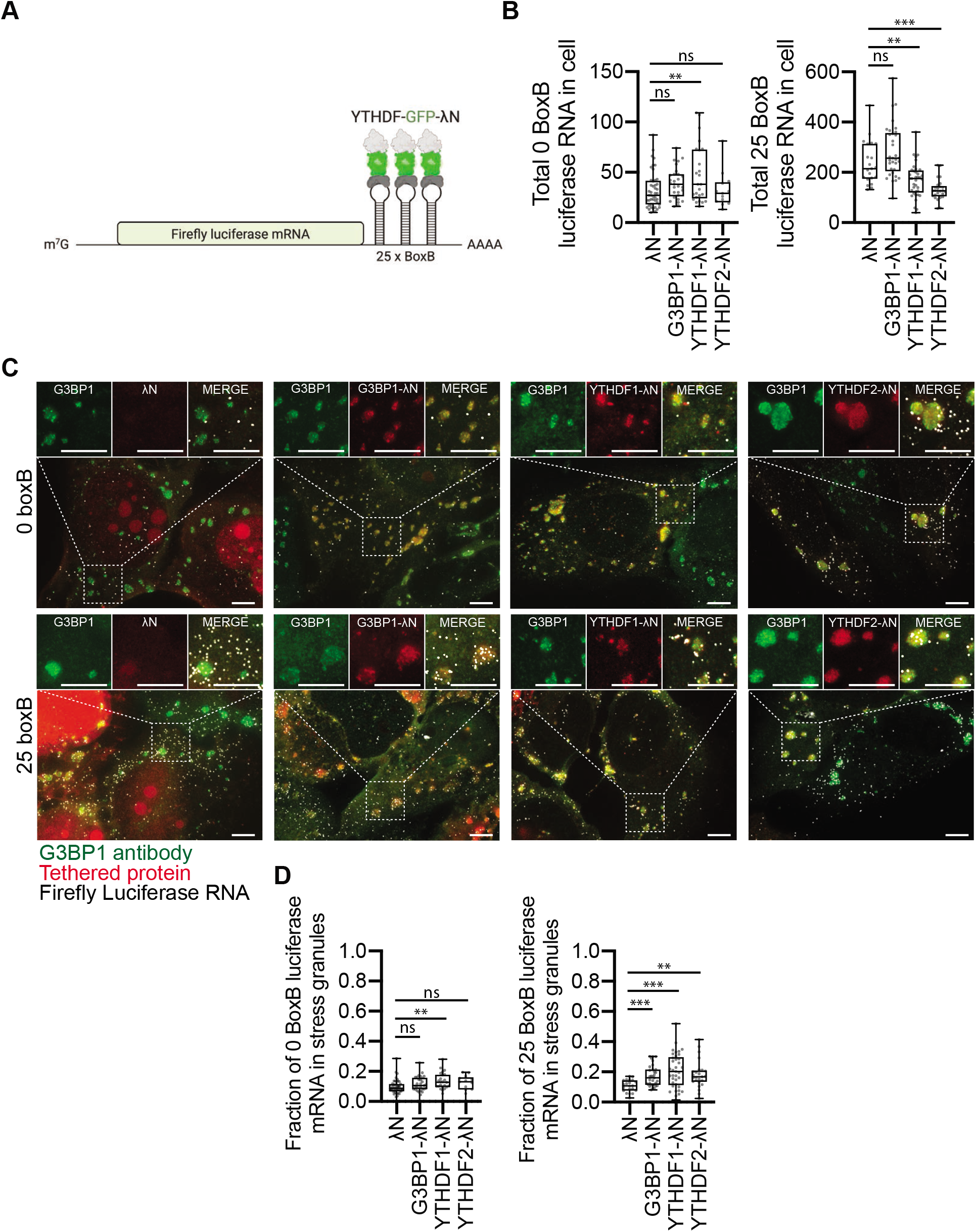
Luciferase RNA is modestly recruited to stress granules when tethered with G3BP1, YTHDF1, or YTHDF2.

(**A**) Cartoon schematic of tethering YTHDF proteins to luciferase mRNA using the BoxB-λN approach. (B) Number of 0 or 25 BoxB luciferase mRNA in U-2 OS cells when expressing λN, G3BP1-GFP-λN, YTHDF1-GFP-λN, and YTHDF2-GFP-λN transgene. U-2 OS cells were stressed for 1 hour with arsenite. Each dot represents a cell. (C) Representative images of arsenite stressed-U-2 OS cells stably expressing 0 or 25 BoxB luciferase mRNA and transiently expressing GFP-λN, G3BP1-GFP-λN, YTHDF1-GFP-λN, and YTHDF2-GFP-λN. Single-molecule FISH probes are illustrated in white, endogenous G3BP antibody staining in green, while tethered proteins are in red. The nuclei are stained with DAPI (blue). The scale bar is 3 μm. (D) Box plot of the fraction of 0 and 25 BoxB luciferase RNA molecules in stress granules in wild-type U-2 OS cells. Each dot represents a cell. **, *, and ns denote P < 0.01, P <0.5, and not significant respectively.

We observed that tethering YTHDF1 or YTHDF2 could increase the recruitment of the 25-BoxB-luciferase reporter mRNA to stress granules from 11% to 21 or 18% for YTHDF1-GFP-λN and YTHDF2-GFP-λN respectively (Figure 3C, D). This effect required the tethering of the YTHDF proteins to the reporter mRNA since no major difference in stress granule recruitment was observed with the 0-BoxB-luciferase reporter control mRNA (Figure 3C, D). In side-by-side experiments with the same cell line, the level of recruitment by YTHDF proteins was similar to the effect of tethering G3BP-GFP-λN (18%). Therefore, these experimental results support the idea that the recruitment of YTHDF proteins to mRNA can increase their partitioning into stress granules, with an approximate contribution similar to the effects of G3BP1.

In summary, we present two experimental approaches indicating that m^6^A modification provides a minimal role in mRNP targeting to stress granules. First, in a direct test of the role of m^6^A modification, we observed that cells deficient in m^6^A modification do not show differences in the partitioning of known m^6^A modified mRNAs into stress granules. However, we provide evidence from multiple regression analysis and tethered function assays that m^6^A modification, presumably by the recruitment of YTHDF proteins, can contribute at least a small effect to mRNA recruitment to stress granules (Figure 2 and 3). By modeling the impact of m^6^A modifications, we estimate that m^6^A will only make a substantial difference in stress granule recruitment for mRNAs that have limited inherrent targeting to stress granules and contain multiple numbers of heavily modified m^6^A sites (Figure 2D, 2F).

Given the limited effect of m^6^A modification on mRNAs partitioning into stress granules, how can we understand mRNA recruitment into stress granules? Our results with m^6^A modification are consistent with the model wherein RNA targeting to granules is a summative effect involving many interactions and no single individual protein-RNA interaction dominates, including m^6^A-YTHDF interaction (Matheny et al., 2021). This model explains why RNA length, which is likely highly correlated with valency, which is the number of interactions the mRNA can make with other RNAs and proteins, is so strongly correlated with enrichment in stress granules (Khong et al., 2017). This model also explains why length correlation with stress granule enrichment is a consistent metric in other cell types with different genes expressed (Khong et al., 2017; Namkoong et al., 2018). We suggest that this summative effect may be a general property of RNA targeting to RNP granules because length bias is also seen in the RNAs that accumulate in P-bodies (Matheny et al., 2019), P-granules (Lee et al., 2019), and BR-Bodies (Al-Husini et al., 2020). Therefore, like stress granules, no single individual protein-RNA interactions will substantially affect RNP granule partitioning.

If the m^6^A modification does not alter RNA partitioning into stress granules, why does the knockdown of multiple YTHDF proteins (Fu & Zhuang, 2020) lead to a decrease in stress granule formation? We suggest the explanation for this observation may be that the IDRs of the YTHDF proteins act as non-specific assembly factors by forming positive interactions with other components of stress granules once the granules are assembled by other specific interactions. This is similar to observations both *in vitro* and in cells that promiscuously interacting IDRs, which can interact with many other proteins, increase the assembly of condensates by shifting the assembly diagram towards increased assembly once the promiscuous IDRs are concentrated in the condensate by specific interactions (Protter et al., 2018). Thus, we suggest that YTHDF proteins, once bound to mRNAs through m^6^A modifications, are recruited into stress granules, and then through promiscuous interactions with other components of the stress granules enhance the assembly of these RNP granules.

## ACKNOWLEDGEMENTS

We like to thank Pedro Batista (NIH) for providing us with both wildtype and knockout mES cells and for providing instructions on how to maintain the cells. We like to thank John Rinn for providing FCS for growing mES cells. Finally, we like to thank Ye Fu and Xiaowei Zhuang (Harvard University) for stimulating and helpful discussions. This work was supported by Banting Postdoctoral Fellowship (A.K.) and Howard Hughes Medical Institute (R.P.)

## MATERIALS AND METHODS

### Multiple linear regression analysis

Differential isoform enrichment was performed as previously described (Khong et al. 2017) (Supplemental Data 1). m^6^A mapped sites and m^6^A ratio were obtained from Xiang *et al*. (2017) and Molinie *et al.*, (2016). The stress granule enrichment dataset was filtered to only consider the most highly expressed isoform for each gene. The m^6^A dataset used Refseq annotation, while our original dataset used Ensembl gene ID’s. In order to assign m^6^A peaks to genes in our stress granule isoform data, we matched Refseq annotations to gene names using pybiomart, counted the number of m^6^A peaks per gene, and merged the two datasets. ORF lengths were also obtained using pybiomart.

Multiple linear regression analysis was performed using the scikit-learn package in python. Multiple models were constructed using ORF length, the number of m^6^A sites, and the combination as features and the observed fraction of molecules within stress granules as a response variable. r^2^ values were obtained to assess the predictive power for each of these models by comparing predicted vs. observed stress granule enrichment. The visualization of our linear model was created by simulating transcripts of varying ORF lengths up to 30kb in size and of varying numbers of m^6^A sites and running these data through our multiple linear regression model.

All visualizations in Figure 2 were created using the matplotlib and seaborn packages in python. All code pertaining to this analysis is contained within the following github repo (which will be released upon manuscript acceptance).

### Cell lines

Wildtype and ΔMETTL3 mES cells were maintained in DMEM supplemented with 15% FCS (ES-009-B, Millipore Sigma), 1X non-essential amino acids (11-140-050, Gibco), 1X penicillin-streptomycin-glutamine (10378016, Thermo Fisher Scientific), 1 mM sodium pyruvate (11360070, Thermo Fisher Scientific), 1X EmbryoMax 2-mercaptoethanol (ES-007-E, Millipore Sigma), 10000 units/mL ESGRO Leukemia Inhibitory Factor (ESG1106, Sigma-Aldrich), 1 μM PD0325901 (S1036, Selleck Chemicals), and 3 μM CHIR99021 (SML0146, Sigma-Aldrich) and maintained at 37°C and 5% O2 on 0.1% gelatin (G6144, Sigma-Aldrich) coated tissue culture plates.

U-2 OS cells were maintained in DMEM supplemented with 10% FBS, 1% penicillin/streptomycin at 37°C and 5% O2. U-2 OS cells stably expressing 0- and 25-BoxB-luciferase RNA was generated by transducing pLenti-EF1-luciferase-blast and pLenti-EF-luciferase-25-BoxB-blast. The plasmids were created by inserting luciferase and luciferase 25-BoxB derived from plasmids described in Matheny et al., (2021) into pLenti-EF1-Blast vector following Gibson assembly. To generate the Luciferase-25-BoxB and Luciferase lentiviral particles, HEK293T cells (T-25 flask; 80% confluent) were co-transfected with either 2.7 μg of either pLenti–EF1–Luciferase-25-BoxB–blast or pLenti–EF1–Luciferase–blast, 870 ng of pVSV-G, 725 ng of pRSV–Rev, and 1.4 μg of pMDLg–pRRE using 20 μl of Lipofectamine 2000. The medium was replaced 6 h post-transfection. The medium was collected at 24 and 48 h post-transfection and filter-sterilized with a 0.45-μm filter. To generate the U 2-OS luciferase and luciferase-25-BoxB stable cell lines, U 2-OS cells (T-25 flask; 80% confluent) were transduced with 1 ml of lentiviral stocks containing 10 μg/ml of Polybrene for 1 h. The medium was then added to the flask. 24 h post-transduction, the cells were reseeded in T-25 flask containing 5 μg/ml of blasticidin selective medium. Single colonies were selected and colonies with an adequate expression of RNA were chosen for this project (20-500 mRNA copies per cell).

### Absolute quantification of m^6^A levels on mRNAs in wildtype and ΔMETTL3 mES cells

Total RNA was isolated from wildtype and ΔMETTL3 mES cells using Trizol reagent (15-596-018, Thermo Fisher Scientific) by following the manufacturer’s protocol. Poly(A) mRNA was then extracted using Dynabeads mRNA purification kit (61006, Thermo Fisher Scientific) by following the manufacturer’s protocol. Absolute quantification of m^6^A levels on poly(A) mRNA was determined using Epiquick m^6^A RNA methylation quantification kit (P-9005-48, EpiGentek) by following the manufacturer’s protocol. Biological replicates were performed to confirm results are reproducible.

### Generation and transfection of GFP-λN, G3BP1-GFP-λN, YTHDF1-GFP-λN, and YTHDF2-GFP-λN plasmids

The creation of GFP-λN and G3BP1-GFP-λN plasmids were previously described in Matheny et al., (2021). YTHDF1 and YTHDF2 were synthesized by Twist Biosciences and subcloned into G3BP-GFP-λN vector by swapping out G3BP1 with YTHDF1 and 2 genes using EcoR1 and BamH1 sites. Inserts were sequenced verified.

### Sequential immunofluorescence and single-molecule FISH

U-2 OS, wildtype and ΔMETTL3 mES cells were seeded on EtOH-sterilized, +0.1% gelatin-coated (mES cells), 18 x 18 mm #1.5 coverslips in six-well tissue culture plates overnight. Cells were stressed by replacing the media with media containing 500 μM NaAsO_2_ and incubating for 1h at 37°C/5% CO_2_, followed by washing with prewarmed 1X PBS and fixing with 500 μl of 4% paraformaldehyde for 10 min at room temperature. After fixation, cells were washed twice with RNase-free 1X PBS (10010049, Thermo Fisher Scientific), permeabilized with 0.1% Triton X-100 in RNase-free 1X PBS for 5 min, and washed once with RNase-free 1X PBS. Coverslips were then stained with primary stress granule antibodies (5 μg/mL mouse α-G3BP primary antibody (ab56574; Abcam) or 1 μg/mL Rabbit α-PABP primary antibody (ab21060; Abcam)) in RNase-free
1X PBS for 60 mins at room temperature, followed by 3 washes with RNase-free 1X PBS, and subsequently stained with respective secondary antibodies (1:1000 goat α-mouse FITC-conjugated secondary antibody (ab6785; Abcam) or 1:1000 donkey anti-rabbit Alexa-fluor 555 antibody (ab150062, Abcam)) in RNase-free 1X PBS for 60 mins at room temperature. Coverslips were then washed 3 times with RNase-free 1X PBS and fixed with 500 μl of 4% paraformaldehyde for 10 min at room temperature.

After antibody staining, single-molecule FISH was performed as previously described with slight changes (Khong et al., 2018). We now make our own buffers instead of using Biosearch Technologies Stellaris buffers (SMF-HB1-10, SMF-WA1-60, and SMF-WB1-20). The recipe for Wash buffer A is 15 mL 20X nuclease-free SSC plus 120 mL nuclease-free water. The recipe for Wash buffer B is 15 mL 20X nuclease-free SSC plus 135 mL nuclease-free water. The recipe for hybridization buffer is 2 mL 50% dextran sulfate, 1 mL deionized formamide, 1 mL 20X nuclease-free SSC, and 6 mL nuclease-free water. All buffers are filter sterilized with syringes and syringe filters. Prior to use, formamide is added to wash buffer A at 10% v/v. Formamide is no longer added prior to use for hybridization buffer. The rest of the protocol for single-molecule FISH is identical to Khong *et al*., (2018).

Single-molecule FISH probes except for the POLR2A (SMF-2006-1, Biosearch Technologies) and firefly luciferase mRNA single-molecule FISH probes (Matheny et al., 2021) are now created using a method as described in Gaspar et al., (2017) (Gaspar et al., 2017). Oligo sequences can be found in Supplemental Table 1.

Imaging parameters are performed as described previously in Khong et al., (2018). Each data point in Figure 1 represents one biological replicate with at least 361 single-molecule FISH spots counted. Each data point Figure 2 represents one cell. Only cells that were adequately expressing the transgene GFP-λN, G3BP-GFP-λN, YTHDF1-GFP-λN, and YTHDF2-GFP-λN were counted (50,000 < Cell Total Cell Fluorescence < 2,000,000 (determined by ImageJ)) in Figure 2. The images were blinded, and the single-molecule FISH spots were manually counted. Total numbers of spots outside and inside stress granules were determined. All images shown in the manuscript are deconvolved and the brightness/contrast adjusted to best indicate data.

---

**Supplemental Table 1.**
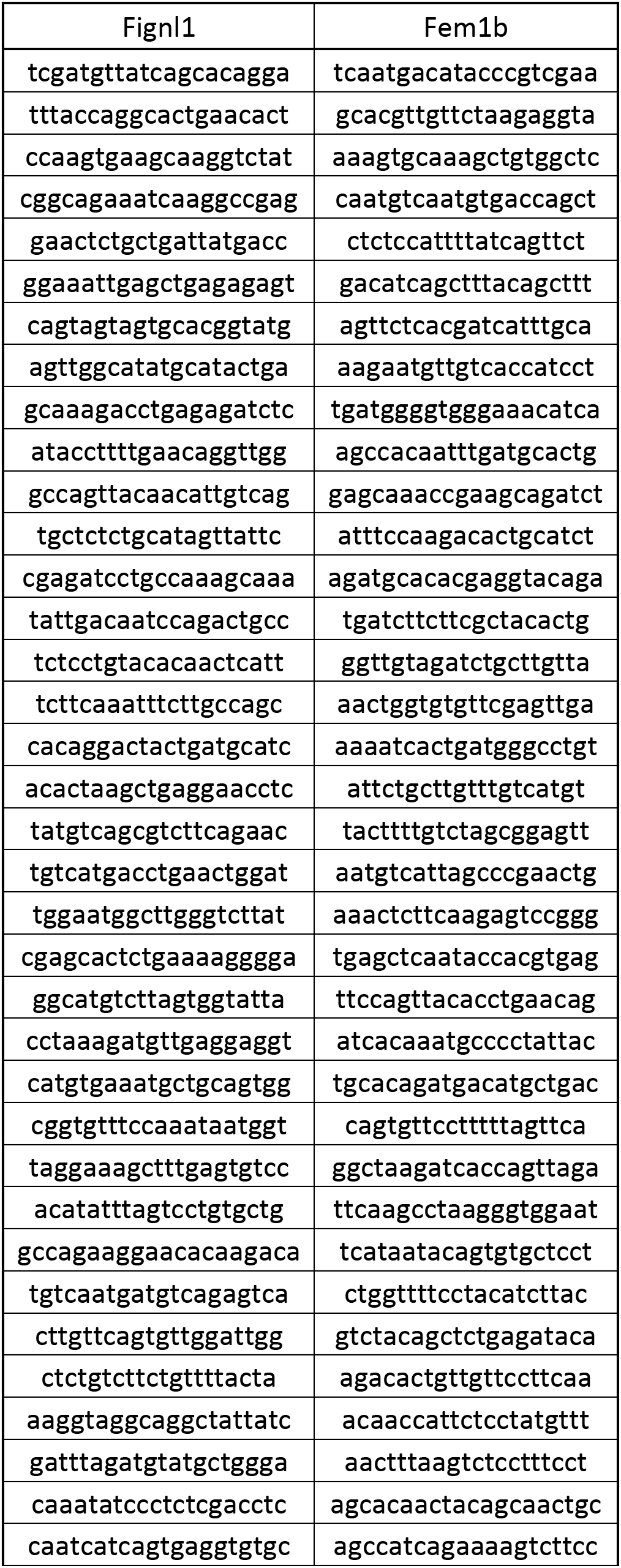
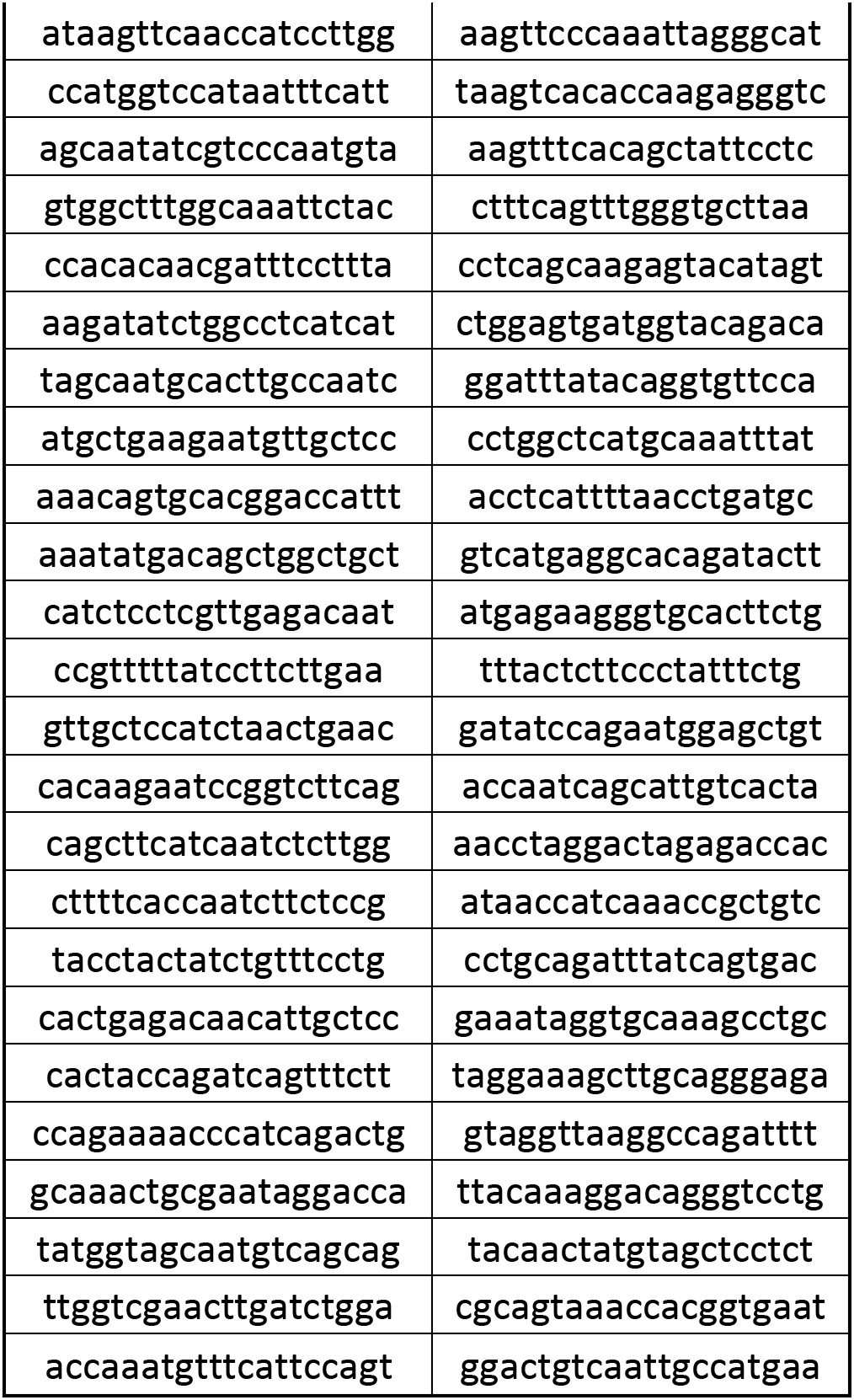
Single-molecule fish probe set sequences.

**Supplemental Figure 1.**
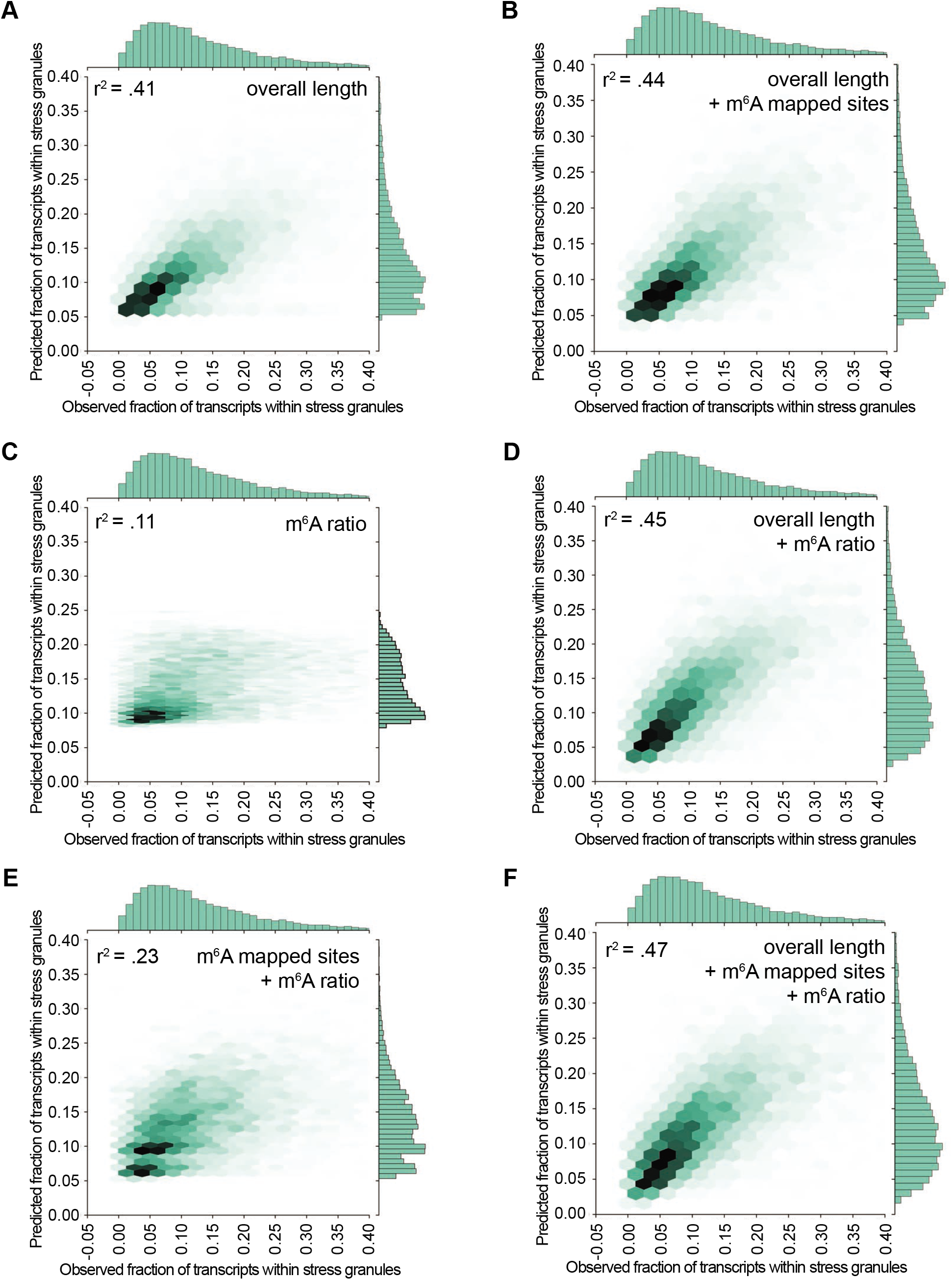
Multiple regression analysis between overall transcript length, m^6^A mapped sites, and m^6^A ratio on RNA enrichment in stress granules. Scatterplot depicting predicted vs observed fraction of transcripts within stress granules with the following metrics: (A) overall length, (B) overall length + m^6^A mapped sites, (C) m^6^A ratio, (D) overall length + m^6^A ratio, (E) m^6^A mapped sites + m^6^A ratio, and (F) overall length + m^6^A mapped sites + m^6^A ratio.

## Notes

### Competing Interest Statement

Roy Parker is a founder and consultant for Faze Medicines.

